# Evolutionary and Functional Implications of Hypervariable Loci Within the Skin Virome

**DOI:** 10.1101/078170

**Authors:** Geoffrey D. Hannigan, Qi Zheng, Jacquelyn S. Meisel, Samuel Minot, Frederick D. Bushman, Elizabeth A. Grice

## Abstract

Localized genomic variability is crucial for the ongoing conflicts between infectious microbes and their hosts. An understanding of evolutionary and adaptive patterns associated with genomic variability will help guide development of vaccines and anti-microbial agents. While most analyses of the human microbiome have focused on taxonomic classification and gene annotation, we investigated genomic variation of skin-associated viral communities. We evaluated patterns of viral genomic variation across 16 healthy human volunteers. HPV and *Staphylococcus* phages contained 106 and 465 regions of diversification, or hypervariable loci, respectively. *Propionibacterium* phage genomes were minimally divergent and contained no hypervariable loci. Genes containing hypervariable loci were involved in functions including host tropism and immune evasion. HPV and *Staphylococcus* phage hypervariable loci were associated with purifying selection. Amino acid substitution patterns were virus dependent, as were predictions of their phenotypic effects. We identified diversity generating retroelements as one likely mechanism driving hypervariability. We validated these findings in an independently collected skin metagenomic sequence dataset, suggesting that these features of skin virome genomic variability are widespread. Our results highlight the genomic variation landscape of the skin virome and provide a foundation for better understanding community viral evolution and the functional implications of genomic diversification of skin viruses.

## INTRODUCTION

Localized genomic modifications are ammunition in the ongoing battle between hosts and infectious agents. The human adaptive immune response relies on localized genomic diversity of antigen receptors to facilitate detection and efficient removal of foreign agents (Borghans, Beltman & De Boer, 2004; Kubinak et al., 2012). Infectious microbes, such as bacteria and viruses, likewise rely on genomic variation to modulate tropism and facilitate immune evasion (e.g. (Malim & Emerman, 2001; Doulatov et al., 2004; Minot et al., 2012; Schillinger et al., 2012; Das et al., 2013; Minot et al., 2013; Guo et al., 2014)). Potential selective benefits of targeted variation in viruses include immune evasion and widening of host tropism (Borghans, Beltman & De Boer, 2004; Kubinak et al., 2012).

Most contemporary low resolution studies of the human microbiome evaluate functional potential through taxonomic classification and whole gene identification (e.g. (Schloss & Handelsman, 2008; Human Microbiome Project Consortium, 2012; Langille et al., 2013; Hannigan et al., 2014; Ly et al., 2014; Norman et al., 2015; Lim et al., 2015; Meisel et al., 2016)). These approaches are usually unable to capture nucleotide variations that affect functionality of proteins encoded in the microbiome, which can be altered by differences in only a few nucleotides. For example, viruses such as *Bordetella* bacteriophages, hepatitis C virus, and others only require short variable regions within a gene to facilitate functional changes in processes including tropism diversity, immune evasion, drug resistance, and adaptation to host auxotrophies (Bacher, Bull & Ellington, 2003; Doulatov et al., 2004; Donaldson et al., 2010; Guan et al., 2012; Shah et al., 2014). Contemporary low-resolution studies also fail to identify genetic cassettes that promote targeted diversity, such as diversity generating retroelements (DGRs). DGRs promote targeted genetic diversification in bacteriophages through error-prone cycles of transcription, reverse transcription, and integration; through this process information encoded in a nonvariable template region is copied in an error-prone fashion into a variable region within a coding sequence (Doulatov et al., 2004; Minot et al., 2012; Schillinger et al., 2012).

Here we investigate skin virome evolution and adaptation by inferring the selective pressure, functional diversity, and substitution patterns associated with targeted hypervariation. We focus on three prominent cutaneous viruses: Human Papillomavirus (HPV), *Propionibacterium* phage, and *Staphylococcus* phage. HPV is associated with the development of skin cancer, especially in immune-suppressed individuals (Vinzón et al., 2014; Wang et al., 2014; Quint et al., 2015). Current vaccine efforts aim to target conserved antigens for broad strain protection—thus a greater understanding of HPV genomic diversity could improve design of vaccines (Schiller & Lowy, 2012; Vinzón et al., 2014). *Staphylococcus* phages can modulate *Staphylococcus* pathogenic gene expression and facilitate transmission of antibiotic resistance (Bae et al., 2006; Varga et al., 2012). *Propionibacterium* phages are associated with *Propionibacterium acnes* (the bacterium associated with acne vulgaris) and have therapeutic potential for treating acne (Marinelli et al., 2012; Hannigan & Grice, 2013). Our findings build upon previous analyses of individual virus genomic variability and provide new insight into phage biology of the cutaneous microbiome.

## MATERIALS & METHODS

### Analysis Details and Availability

All associated source code and explanatory README files are available for review at the following GitHub respository: https://github.com/Microbiology/ViromeVarScripts.

### Data Acquisition & Quality Control

The primary skin virome dataset was acquired from SRA accession: SRP049645 (Hannigan et al., 2015). Sequences from samples collected at time point two and three were downloaded. Assembled contigs were downloaded from the FigShare source (doi: 10.6084/m9.figshare.1281248).

The secondary dataset was obtained from Oh *et al* (SRA BioProject: 46333) (Oh et al., 2014). Retroauricular crease samples were downloaded from the NCBI SRA BioProject: 46333. Sequences were trimmed by quality score using a cutoff of 33 and the FastX toolkit (version 0.0.14). Remaining sequences with similarity to the human genome were removed using the standalone DeconSeq toolkit (version 0.4.3) (Schmieder & Edwards, 2011).

### Contig Assembly and Taxonomic Identification

Contigs from the primary dataset were obtained from the published Figshare source (doi: 10.6084/m9.figshare.1281248). These contigs were assembled using the Ray metagenomic assembly software (v2.3.1) (Boisvert et al., 2012). Contigs from the Oh *et al* dataset were also assembled using the Ray assembler. Within each dataset, sequences from all samples were combined prior to assembly to facilitate the most complete contig assembly. Contig coverage was determined by aligning sequences back to the contigs with the bowtie2 toolkit (v2.1.0; seed substring length of 25 and one mismatch allowed in alignment) (Langmead & Salzberg, 2012). Quantification of reads mapping back to contigs was obtained by parsing bowtie2 output using Perl and BASH scripts as presented in the supplemental source code. Coverage was calculated using the number of reads mapping to each contig. The blastn program from the NCBI Blast+ toolkit (version 2.2.0) was used to determine similarity of contigs to virus reference genomes (Camacho et al., 2009). Contigs were blasted against a previously described virus-specific genome reference database, which is a subset of the EMBL reference genome database (UniProt Consortium, 2014; Hannigan et al., 2015). A similarity threshold of e-value < 10^−3^ was used, and sequences with multiple potential identities were resolved by using only hits with the lowest e-values. Although this was the minimum threshold, the contigs of interest exhibited e-values much lower than 10^−3^.

### Phylogenetic Analysis

We constructed phylogenies using the L1 capsid gene for HPV (Ma et al., 2014) and the large terminase subunit for the *Staphylococcus* and *Propionibacterium* phages (Gutiérrez et al., 2013; Ma et al., 2014) as phylogenetic marker genes. For reference, we used the PAVE reference L1 genes (<pave.niaid.nih.gov>, accessed 2015-06-03) (Van Doorslaer et al., 2013). The large terminase subunit references for *Staphylococcus* and *Propionibacterium* phages were from the NCBI gene sequence database (*Staphylococcus* phage: accessed 2015-09-14, <u>search term</u>: *((phage terminase large subunit staphylococcus)) AND "viruses"[porgn:_txid10239] NOT "ORF" NOT "hypothetical" NOT "putative"*; *Propionibacterium* phage: accessed 2015-09-15, <u>search term</u>: *((phage terminase large subunit propionibacterium)) AND "viruses"[porgn:_txid10239]*). To extract the phylogenetic marker genes from the virome contigs, we determined which open reading frames (ORFs) matched the reference genes by nucleotide similarity (nucleotide blast, evalue 1e-10). Only ORFs longer than 1.2 kb were included in the analysis. Contig and reference marker genes were aligned using the Smith-Waterman algorithm and 1000 iterations as implemented by the mafft aligner (v7.221) (Katoh & Standley, 2013). Phylogeny was constructed using RAxML (version 8.1.21) (Stamatakis, 2014). The phylogenetic tree was visualized using Figtree (Rambaut).

### Identification of Hypervariable Loci

The bowtie2 read alignments were formatted (e.g. conversion from binary to ASCII format) using the Samtools toolkit (v1.1), and then SNPs were called using VarScan (v2.3.7) (Li et al., 2009; Koboldt et al., 2012). The ‘pileup2snp’ program from VarScan was used with a minimum minor allele frequency threshold of 1%, a read depth of 8, and a minimum of two supporting reads for variant calls. Indels were excluded.

To identify hypervariable loci, we used a geometric distribution based statistic approach as described previously (Zheng et al., 2010), which has the advantage of avoiding boundary difficulties and variations within contigs compared to similar methods such as sliding window searches. We first fit the observed nucleotide distance between SNPs to independent geometric models for each contig, and then identified runs of consecutive SNPs that were closer together than expected by chance using a 95% confidence interval.

### Protein Family Domain Identification Within Hypervariable Loci ORFs

Protein family domains were identified in ORFs that contained hypervariable loci. The subset of translated virus ORFs that contained hypervariable loci were aligned to the standard Pfam protein family domain database using hmmscan within the HMMer toolkit (version 3.1) and GA gathering bit score thresholds (Finn, Clements & Eddy, 2011).

### Prediction of Single Amino Acid Variant Effect on Phenotype

The SuSPect algorithm was employed to predict the likelihood of SNP-associated single amino acid variants (SAV) impacting phenotype. We used SuSPect to create a matrix of likelihood scores for every possible SAV at every position in the ORFs that contained hypervariable loci. This matrix was used as a reference to quantify the likelihood of each hypervariable loci SNP to impact the resulting phenotype. The significance of the score differences between viruses was calculated using a Wilcoxon rank-sum test.

### Evolutionary Pressure of Hypervariable Loci and Virus Genomes

We assessed the evolutionary pressure of a gene using the ***pN***/***pS*** ratio as in **Formula 1**, where M_N_ and M_S_ represent the observed number of non-synonymous and synonymous SNPs, respectively. These values were normalized by the total number of possible non-synonymous or synonymous substitutions (N_i_ and S_i,_ respectively), in order to avoid potential codon usage bias. Furthermore, to normalize for sequence coverage of the SNPs and prevent extreme values, a pseudocount value was added to the M_N_ and M_S_ values, which was defined as half of the square root of the median sequence coverage of SNPs within the region of interest (C_M_). The pseudocount approach was used to prevent infinite and illegal values when M_N_ or M_S_ had zero values, thus allowing consideration of otherwise ignored data points.

This analysis is similar to the ***dN***/***dS*** calculations often performed to estimate degrees of natural selection among genomes (Nishida, Frith & Nakai, 2009; Schloissnig et al., 2013). It is important to note however that such an analytical approach would be inappropriate for this type of sample set because the nucleotide variations are not assignable to isolated strains, which prevents haplotype identification that is a necessary component of ***dN/dS*** calculations. The ***pN**/**pS*** calculation does not assume haplotypes, and is therefore appropriate for metagenomic datasets.

To estimate the selective pressure on hypervariable loci, the locations of the hypervariable loci were extracted, along with their immediately adjacent regions, using a Perl script as presented in the supplemental source code. These positions were used with the contig sequences, SNP call data, and ***pN**/**pS*** calculator to estimate their selective pressure. Hypervariable regions outside of coding regions were not considered.

Calculation of the overall selective pressure on virus contigs was performed in a similar approach to the hypervariable loci selective pressure. Predicted ORFs were first extracted from the contigs using the Glimmer3 toolkit (v3.02) (Delcher et al., 2007). The predicted ORFs, along with the contig sequences, SNP profile, and ***pN**/**pS*** calculator were used to calculate the overall selective pressure on each gene within each contig. The distributions of selective pressures observed for each gene were observed as categorized by virus type.

### Amino Acid Frequency, Charge, and Polarity

Amino acid abundance profiles were calculated while correcting for the random probability of that substitution. More specifically, each value was weighted for the number of nucleotides that result in the same amino acid as weighted value = *((number of nucleotide substitutions resulting in same amino acid) / 3)^−1^*. Relative abundance was calculated as the sum of the corrected frequencies. Charge and polarity were determined using a simple table of known amino acid properties. Differences in profiles between viruses were calculated using a chi-square test.

### Diversity Generating Retroelement Identification

We identified potential diversity generating retroelements (DGRs) by collecting assembled contigs that contained open reading frames similar to known reverse transcriptase genes, and a duplicated nucleotide region less than 150bp in length. Reverse transcriptase open reading frames were identified using blastx (e-value < 10^−5^) and the Uniprot reference reverse transcriptase sequences (http://www.uniprot.org/uniprot/?sort=score&desc=&compress=yes&query=%22reverse%20transcriptase%22%20(phage%20OR%20virus)&fil=&format=fasta&force=yes). Repeat regions were identified by comparing each contig to itself with tblastx (e-value < 10^−50^) and were filtered using customt scripts to remove duplicates and regions longer than 150bp. DGR candidates were removed if they contained no hypervariable loci or if the variable region was no within a predicted open reading frame.

Diversity generating retroelements were visualized in the Integrated Genomic Viewer using the DGR cassettes and bowtie2 aligned sequences described above. The linkage disequilibrium was calculated using a custom perl script for formatting and the “LDheatmap” and “genetics” R packages for analysis and visualization (Shin et al., 2006; Warnes et al.).

### Comparison of Primary Analysis to Validation Dataset

Near identical contigs were identified between the primary and secondary validation dataset by aligning the two individually assembled contigs to each other using bowtie2, with a specified seed length of 25 and up to one seed mismatch. Sequences from our primary dataset and the Oh *et al* dataset were aligned to the near identical contigs. These alignments were used to identify shared SNP locations between our dataset and the Oh *et al* dataset. We quantified shared SNP location as percent of our primary analysis SNPs whose location was identical to those of SNPs in the secondary validation dataset. As a control, we compared these results to a simulated dataset where SNP position was randomly assigned. The example SNP alignment over the circular contig was generated using Genious (Kearse et al., 2012).

## RESULTS

### Diversity of Skin Viruses

We evaluated genomic variability associated with dsDNA skin viruses using a previously published human skin virome metagenomic dataset, consisting of 260,714,906 high quality sequences assembled into > 76,000 contigs from 16 individuals (SRA Accession: SRP049645) (Hannigan et al., 2015). We relied on database virus annotation to identify contigs in our dataset with the highest confidence matches to reference genomes. Because greater sequencing coverage allows for more refined detection of variable nucleotides (Schloissnig et al., 2013), we focused our analysis on *de novo* assembled contigs with coverage greater than 10X. Nearly all contigs that met these criteria were identified as *Propionibacterium* phages (contig count = 45), *Staphylococcus* phages (contig count = 319), and Human Papillomaviruses (HPVs; contig count = 56; **Figure 1A**). A minority of the contigs were identified as *Pseudomonas* phages or *Enterobacteria* phages, but were excluded from the analysis because their annotations were lower confidence and contig representation was minimal (**Figure 1A**).

**Figure 1.**
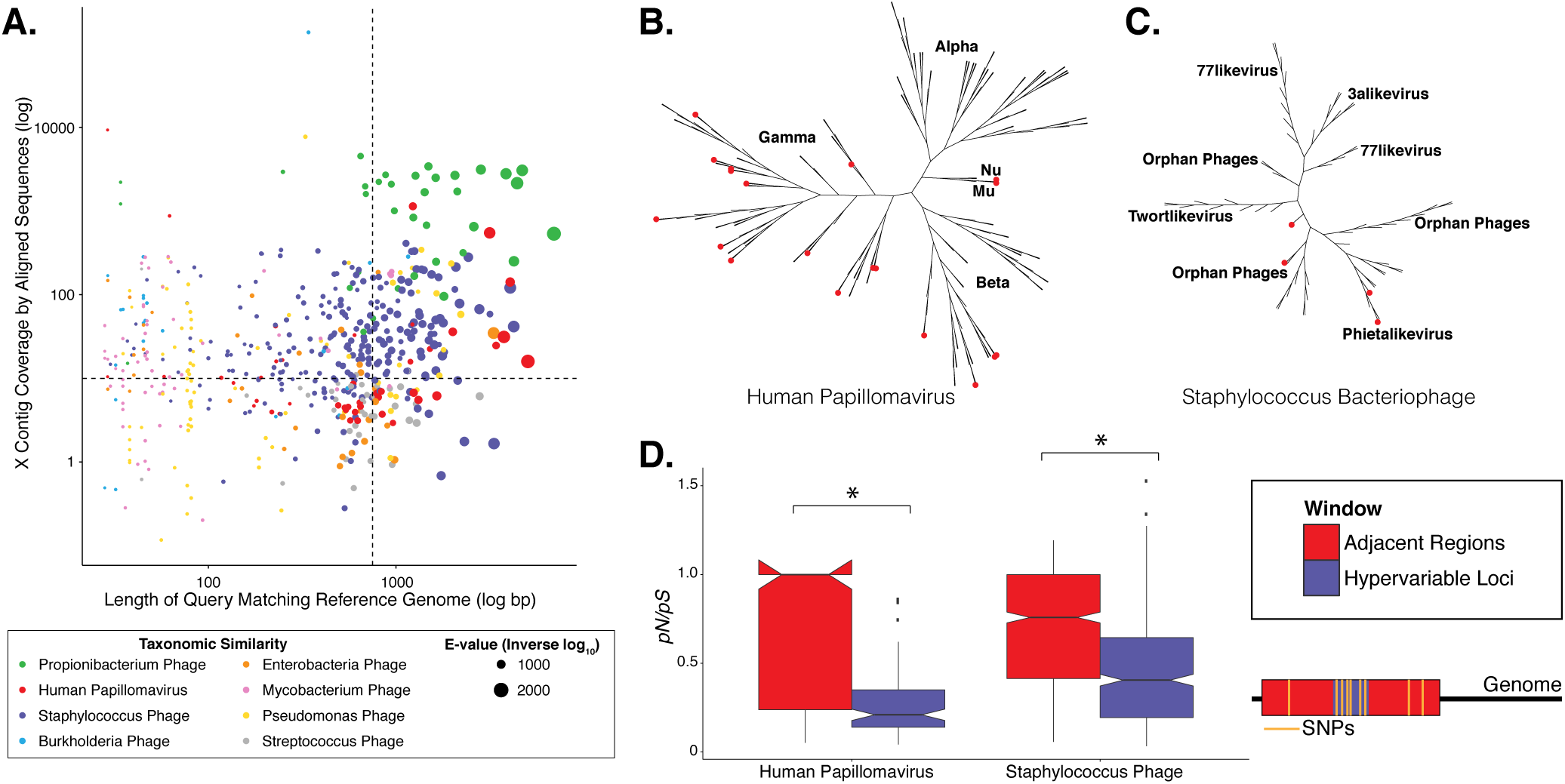
Phylogenetic & Evolutionary Characteristics of Skin Virome Hypervariable Loci A) Scatter plot depicting the candidate contigs considered for analysis in this study. Each point is a contig that mapped to a reference virus genome. The x-axis shows the length (log_10_ scale) of the contig subsection that mapped to the reference genome. The y-axis shows the overall coverage of the contig, as a quantification of sequences aligning to the contig. The color highlights the reference virus genome that the contig was most similar to, and the size depicts the e-value (inverse log_10_) associated with the contig-reference match. The horizontal dashed line marks the threshold of 10X coverage, and the vertical dashed line marks the 750 bp length threshold. **B)** Phylogenetic tree of skin virome HPVs and **(C)** *Staphylococcus* phages, structured onto a standard phylogenetic tree using reference genomes. HPV phylogeny was based on the L1 major capsid gene and *Staphylococcus* phage phylogeny was based on the large terminase subunit. Contigs from this study are highlighted as orange dots, and genera are labeled with text. Phylogenetic lengths were normalized to ranks to facilitate visualization. **D)** Box plots depicting the evolutionary pressure of HPVs (left) and *Staphylococcus* bacteriophages (right) at the hypervariable loci (blue) and the regions immediately adjacent to the hypervariable loci (red). Adjacent regions were calculated as being twice the length of the hypervariable loci (see visualization to the right). The hypervariable locus and adjacent region (combination of both sides) from each sample were evaluated for evolutionary pressure (y-axis) using SNPs (pink lines in right illustration). Asterisk (*) indicates a statistically significant difference (p < 0.01). Notched boxplots were created using ggplot and show the median (center line), the inter-quartile range (IQR; upper and lower boxes), the highest and lowest value within 1.5 * IQR (whiskers), and the notch which provides an approximate 95% confidence interval as defined by 1.05 * IQR / sqrt(n).

We evaluated the diversity of the skin viruses by constructing phylogenic trees based on conserved viral genes. Only fully assembled ORFs were considered of > 1.5 kb for HPV, and > 0.15 kb for *Staphylococcus* and *Propionibacterium* phages. Similar to previous studies, we used the L1 major capsid gene to classify HPV strains (de Villiers et al., 2004; Ma et al., 2014). The terminase large subunit gene was extracted from *Staphylococcus* phage contigs to construct phylogeny as described previously (Gutiérrez et al., 2013; Ma et al., 2014). Because this gene is used for phylogeny of a variety of phages, we attempted to construct *Propionibacterium* phage phylogeny in a similar manner (Ganz et al., 2014; Li et al., 2014), but were ultimately unsuccessful due to the lack of a full-length *de novo* assembled reference genes in the dataset.

Most skin HPVs were identified as gamma HPVs, the prototypical cutaneous HPV class (**Figure 1B**) (Mistry, Wibom & Evander, 2008). A minority of contigs were identified as beta and Mu/Nu HPVs, and none were identified as alpha HPVs. This is consistent with data from the Human Microbiome Project cohort (Ma et al., 2014).

Fewer *Staphylococcus* phage marker genes were identified, compared to HPVs, likely because *Staphylococcus* phage genomes are orders of magnitude longer than HPV genomes, thereby decreasing the probability that contigs covered the entire genome. Because multiple displacement amplification (MDA) was not used to create this dataset, there is no MDA-associated bias toward small circular genomes. The *Staphylococcus* phage contigs belonged to the Phietalikevirus genus and orphan virus groups (those that have not yet been classified) (**Figure 1C**).

### Hypervariable Loci Within the Skin Virome

We implemented a geometric distribution-based approach to identify regions of high genomic diversity, as in (Zheng et al., 2010). Regions within each contig that contained a significantly higher frequency of SNPs over the stochastic background were identified as viral hypervariable loci. HPVs and *Staphylococcus* phages maintained 106 and 465 hypervariable loci, respectively. We were unable to detect hypervariable loci in the *Propionibacterium* phage population.

To determine the virus protein family domains hosting hypervariable loci, we used the Hidden Markov Model (HMM) analysis implemented by HMMer (Finn, Clements & Eddy, 2011). Hypervariable loci-containing HPV genes include E6, E2, and E1 genes, which are associated with infectious gene expression, and the L1 major capsid gene, which is involved in tropism and host immune evasion (**Table S1**). The L1 major capsid protein is also a target in contemporary, widely used HPV vaccines (Schiller & Lowy, 2012). Hypervariable loci were detected in a variety of *Staphylococcus* phage genes with predicted functions related to tropism, host immune evasion, and utilization of host resources (**Table S2**).

### Selective Pressures on Hypervariable Loci

We evaluated the selective pressures on virus genes by calculating the ***pN***/***pS*** ratio of non-synonymous SNPs (***pN***) to synonymous SNPs (***pS***) within each virus taxa (Schloissnig et al., 2013). This was used as an alternative to ***dN/dS*** because ***dN/dS*** assumes haplotype information which cannot be fulfilled by metagenomic data (Schloissnig et al., 2013). In the ***pN***/***pS*** calculation, neutral evolution is defined as an equal frequency of synonymous and non-synonymous polymorphisms. Selective pressure favors non-synonymous mutations, resulting in increased ***pN***/***pS*** ratios. Purifying selection has the opposite effect. Because the existing model (Schloissnig et al., 2013) is susceptible to stochastic effects and extreme outliers (e.g. division by zero when ***pS*** = 0), we added a pseudocount correction (**Formula 1**).

We determined whether hypervariable loci are in fact loci of focused selective pressure by comparing ***pN***/***pS*** values of the loci to the adjacent genomic regions. ***pN***/***pS*** values of hypervariable loci were significantly lower than adjacent regions in both HPV (median: adjacent = 1.0, hypervariable loci = 0.21; p-value = 3.4e-17) and *Staphylococcus* phage (median: adjacent = 0.76, hypervariable loci = 0.41; p-value = 1.8e-40) genomes, suggesting purifying selection and a propensity to maintain existing protein sequences (**Figure 1D-E**). HPV hypervariable loci were under significantly greater purifying selection than those of *Staphylococcus* phages (median: HPV = 0.21, *Staphylococcus* Phage = 0.41; p-value = 4.64e-9) (**Figure S1**).

To evaluate whether the observed selective pressure in HPV and *Staphylococcus* virus communities is genome-wide or localized to hypervariable loci, we quantified the selective pressure on each virus’ genome by calculating the overall ***pN***/***pS*** ratio including hypervariable loci and non-hypervariable loci SNPs. We observed nearly neutral pressure across HPVs and *Staphylococcus phages* that mirrored pressures to those observed in the regions adjacent to the hypervariable loci (median: HPV = 1.0, *Staphylococcus phage* = 0.81, p-value = 3.2e-5) (**Figure S2**).

### Functional Implications of Targeted Substitutions Within Hypervariable Loci

In order to evaluate the specific nucleotide changes occurring at hypervariable loci, as well as to evaluate the implications of specific nucleotide polymorphisms, we quantified the frequency of individual nucleotide substitutions within hypervariable loci. A>C and T>C substitutions were most frequent in HPV hypervariable loci (**Figure 2A**). *Staphylococcus* phages exhibited a significantly different substitution profile (p-value = 0.00018, Chi-Square test), with the most common substitutions being A>G and G>A transitions (**Figure 2B**). HPV and *Staphylococcus* phage substitutions were more likely to be transitions, with a transition/transversion (ti/tv) ratio of 3.25 and 2.02, respectively.

**Figure 2.**
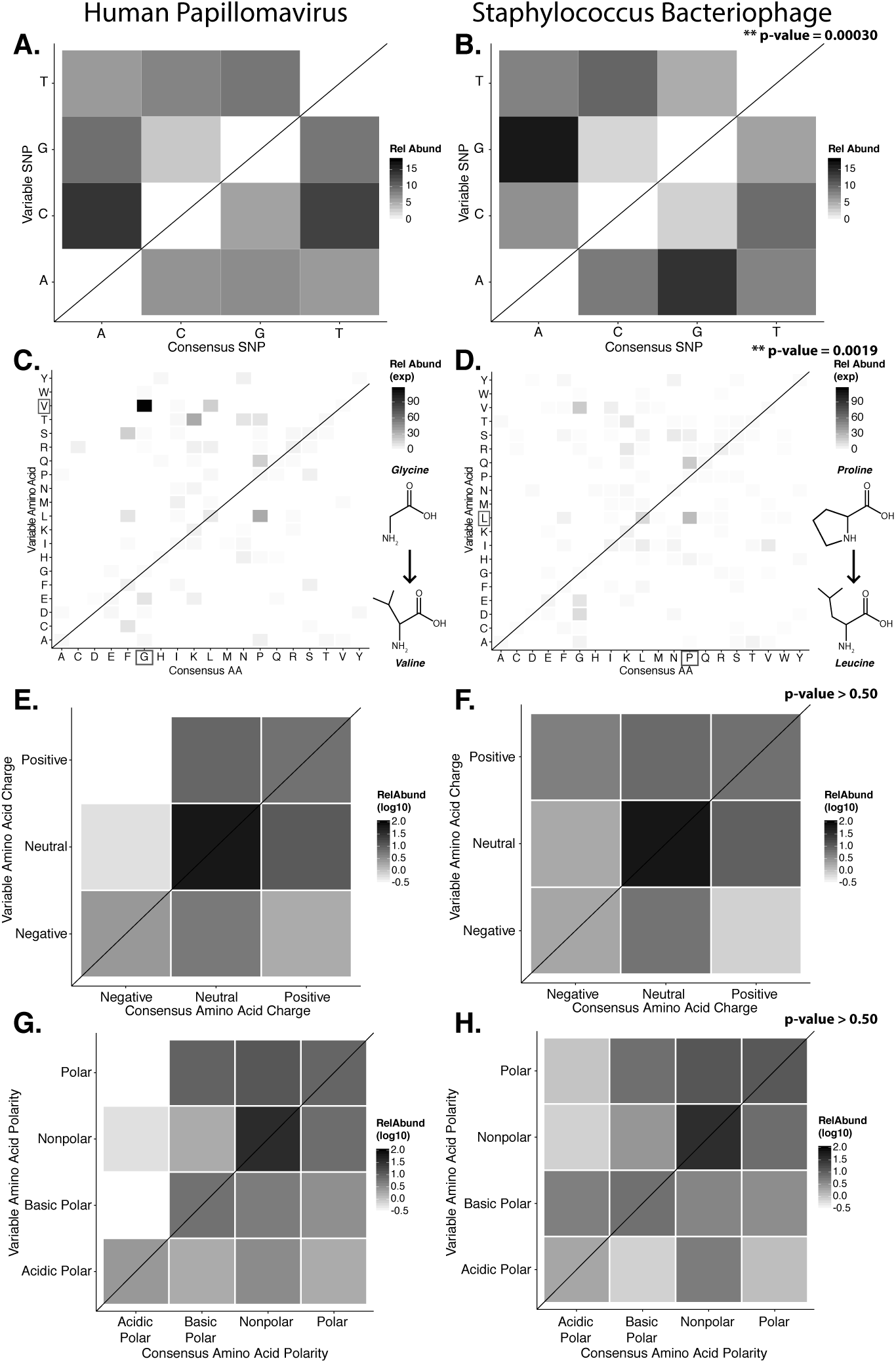
Nucleotide and Amino Acid Substitution Patterns within Viral Hypervariable Loci Heat maps portraying the counts of every possible nucleotide substitution for each SNP found within **(A)** HPV and **(B)** *Staphylococcus* phage hypervariable loci. Tile color weight corresponds to the relative abundance of SNP substitution counts. The diagonal line highlights the panels associated with no substitution. The substitution patterns of amino acids at each SNP are also shown with exponential transformation **(C,D)** An illustration of the major amino acid substitutions are provided beneath the legends as a reference. Amino acid charge **(E,F)** and polarity with acidity **(G,H)** are shown with log_10_ transformation. The absence of a basic or acidic polar identifier indicates the amino acid is polar but neutral. The HPV substitution profiles are found in the left column and the *Staphylococcus* phage profiles are found on the right. Chi-Square significance p-value, comparing variation profiles between the viruses in each row (i.e. A and B), is shown in the upper right corner of the associated *Staphylococcus* phage variation profile. The most frequently substituted amino acid pairs are highlighted with a box around the amino acid letters.

We predicted how hypervariable loci SNPs might affect protein functionality by evaluating patterns of the amino acid substitutions while correcting for the random chance that the substitution will occur. The most frequent non-synonymous amino acid substitution in HPVs was glycine (consensus amino acid) to valine (variant amino acid, **Figure 2C**). While these amino acids are (nonpolar and hydrophobic), glycine is less hydrophobic than valine. The most frequent non-synonymous amino acid substitution in *Staphylococcus* phages was proline to leucine (**Figure 2D**), a substitution between a non-polar cyclic amino acid and an aliphatic straight chain amino acid. Profiles of amino acid substitution were significantly different between HPVs and *Staphylococcus* phages (p-value = 0.0021; Chi-square test).

Amino acid polarity and charge were largely maintained in HPV hypervariable loci (**Figure 2E, G**). In instances of altered charge, visual inspection suggests the most frequent changes were from neutral to positive or negative charge, or positive to neutral charge. Consensus acidic polar residues were not associated with polymorphisms. *Staphylococcus* phage community hypervariable loci appeared to be under weaker substitution selection, with a greater diversity in amino acid charge and polarity (**Figure 2F, H**) compared to HPV. Patterns of substitution charge and polarity were not significant (p-value > 0.5; Chi-square test) when comparing the entire HPV to *Staphylococcus* phage substitution profiles.

We reinforced the observed functional implications of hypervariable loci by predicting the effects of their associated single amino acid variants (SAVs) on gene phenotype using the support vector machine algorithm implemented in SuSPect (Yates et al., 2014). This method assigns a deleterious score to each hypervariable loci SNP-associated SAV, with 0 representing a neutral SAV and 100 representing a SAV with high likelihood to impact phenotype. These scores are based on the predicted impact of the single amino acid variant on the tertiary and secondary structure of the resulting protein, the location of the SAV within the resulting protein (surface vs core), and whether the SAV has previously been associated with altered protein-protein interactions. Both *Staphylococcus* phages and HPVs have an abundance of SNPs associated with SAVs predicted to be deleterious (deleterious scores approaching 100) (**Figure 3**). The HPV SNPs were predicted to be significantly more likely to impact phenotype than the *Staphylococcus* phage SNPs (median: HPV = 45, *Staphylococcus* phage = 17; p-value < 2.2e-16), suggesting that SNPs impact functionality differently between viruses.

**Figure 3.**
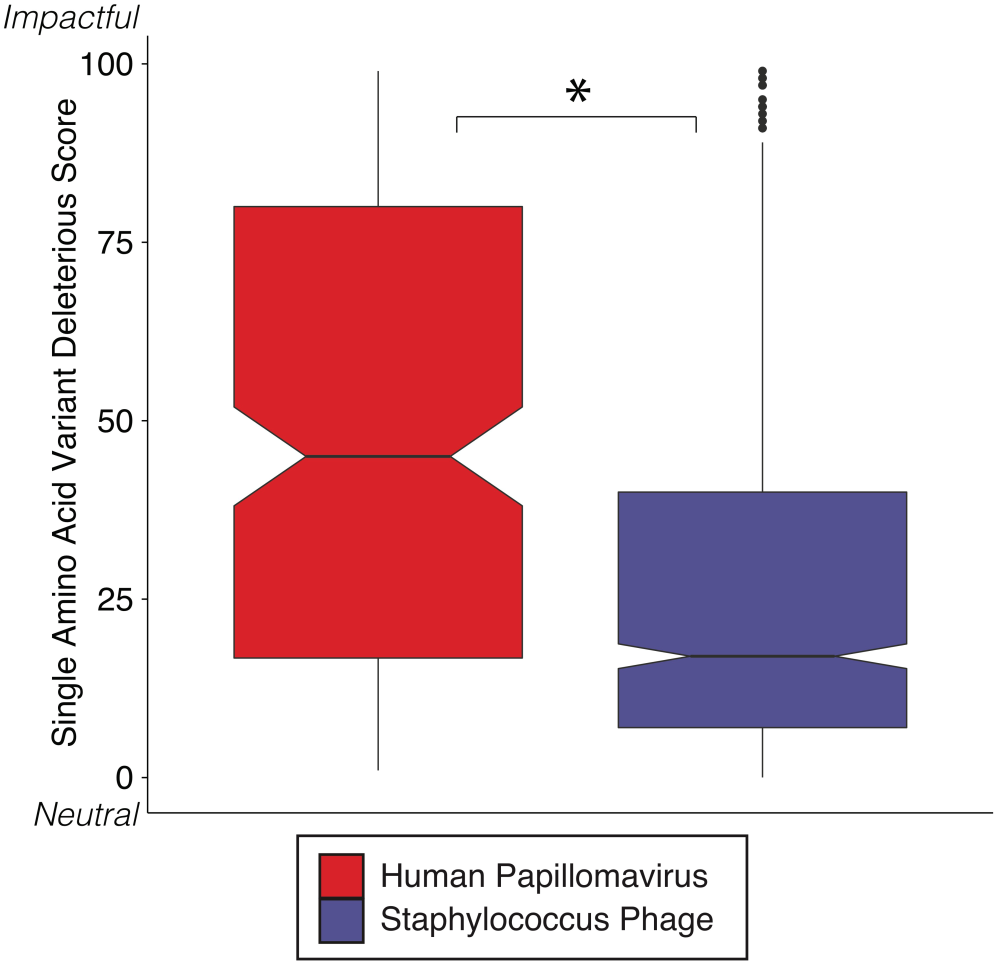
SVM Predicted Impact of Hypervariable Loci on Phenotype Notched boxplot of deleterious scores in Human Papillomavirus (red) and *Staphylococcus* phage (blue) genomes. A low deleterious score indicates a predicted neutral phenotypic effect, while a high score indicates a predicted strong phenotypic effect. Asterisk (*) indicates significant difference by Wilcoxon rank-sum test (p < 1e15). Boxplot parameters as described in figure 1.

### Diversity Generating Retroelements as a Mechanism for Targeted Hypervariability

Diversity generating retroelements (DGRs) are a genetic system used by bacteriophages (as well as bacteria and archea) to promote targeted hypervariability in genes (Doulatov et al., 2004). While DGRs are complex and consist of many components, at their most basic they can be identified as elements consisting of a reverse transcriptase gene and a repeated nucleotide sequence of length < 150 bp that is found in two separate locations of the genome (Doulatov et al., 2004; Minot et al., 2012; Schillinger et al., 2012), termed the template region and the variable region. The template region is transcribed, then reverse transcribed in an error-prone fashion. The resulting cDNA is then integrated into the variable region, introducing base substitutions. Targeted hypervariation impacts functions including broadened host cell tropism by mutagenizing a phage tail fiber gene (Doulatov et al., 2004).

We thus sought to identify candidate DGR cassettes within our viral contigs. We defined the candidate cassettes as pairs of non-overlapping regions with similar nucleotide sequences (tblastx of contigs against themselves, e-value < 10e-50) and co-localized on a contig containing a predicted virus/phage reverse transcriptase gene. We only considered cassettes that were located within a predicted viral gene, contained at least one hypervariable locus in their variable region, and exhibited truly random variation (different between reads). Based on these criteria, we identified one *Staphylococcus* phage DGR candidate that contained hypervariable loci.

For the *Staphylococcus* phage DGR candidate with hypervariable loci, we calculated the linkage disequilibrium associated with the variable nucleotide positions to infer whether the DGR was active or inactive (e.g. an evolutionary artifact). The DGR cassette had unlinked nucleotide variation, which was supported by low levels of linkage disequilibrium (R^2^) in the variable region (**Figure 4**). In this cassette, the template region has less frequent blocks of linkage equilibrium (unlinked variants) while the variable region was associated with greater linkage equilibrium. Together this suggests the observed *Staphylococcus* phage is active. The variable region was associated with a gene of unknown function.

**Figure 4.**
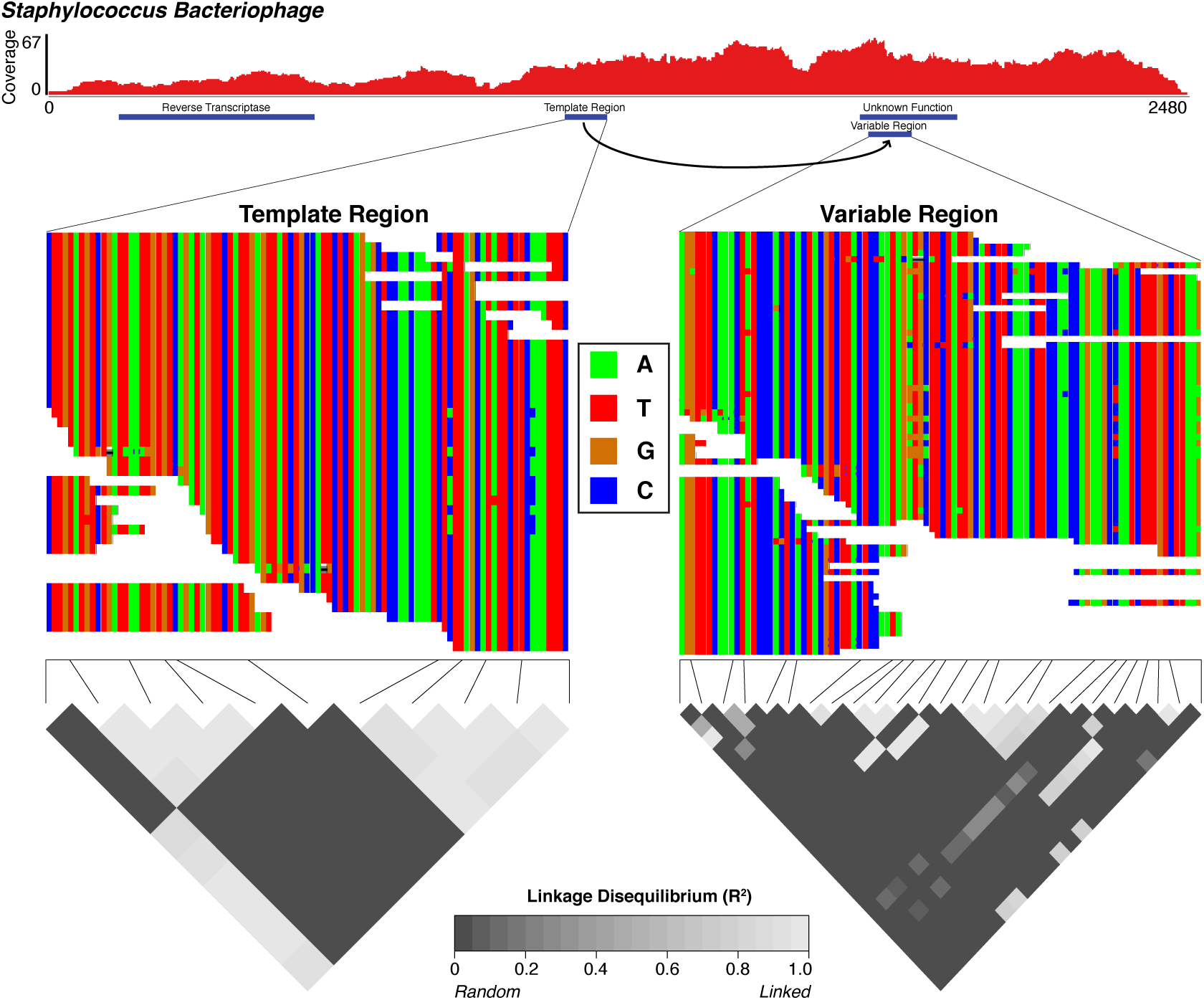
The Diversity Generating Retroelement as a Mechanism for Targeted Nucleotide Variation Alignment illustrating a putative diversity generating retroelement in *Staphylococcus* phage. **Top Panel**) Sashimi plot of sequence coverage across the contig. Coverage ranges from 0-67 X. Below the coverage is a map of the relevant genes predicted within the contig. **Middle Panel**) Sequence alignment of the diversity generating retroelement template region (left) and variable region (right). **Bottom Panel**) Linkage disequilibrium heatmap for the template and variable region. Panels compare variable nucleotides to each other and darker tiles indicate decreased linkage disequilibrium correlation (R^2^; unlinked nucleotide variability).

### Skin Virome Variability Patterns and SNP Locations Are Reproducible Across Different Datasets

We repeated our analyses in a separate, independently collected dataset from another research group (SRA BioProject: 46333) (Oh et al., 2014) to determine the generalizability of our findings. We analyzed metagenomic sequence data of skin specimens that were collected from the retroauricular crease without initial purification of virus like particles. Consistent with our primary analysis, *Staphylococcus* and *Propionibacterium* phages were identified as having the highest coverage and similarity to reference genomes (**Figure 5A**). *Pseudomonas* phages were identified but were in the minority and had low coverage and similarity to reference genomes. HPV was not identified as a major virus in our analysis of the retroauricular crease; however, molluscum contagiosum virus, a poxvirus that causes cutaneous growths that become severe in immunocmpromised states, was present in high relative abundance in agreement with the original published findings (Oh et al., 2014).

**Figure 5.**
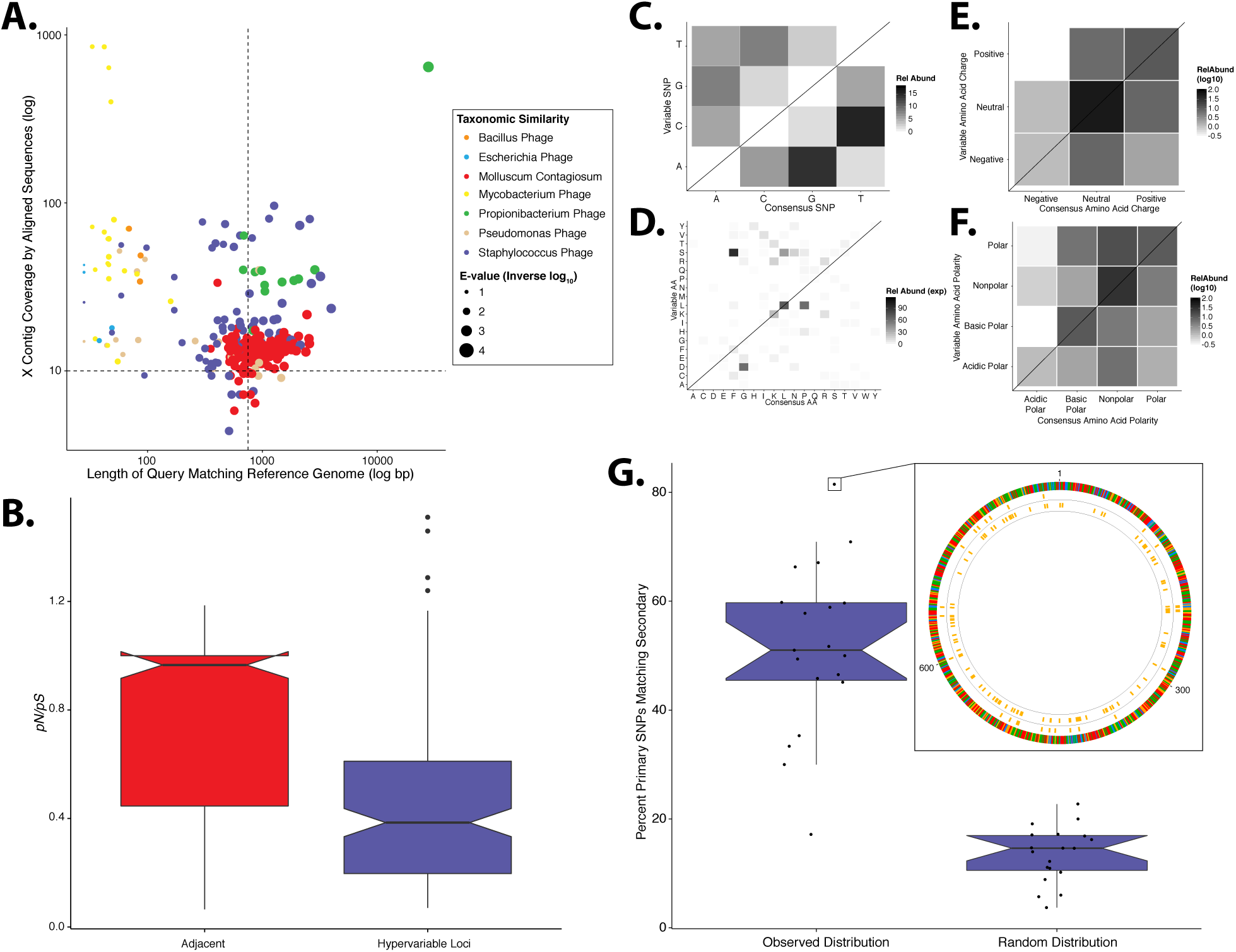
Validation of Study Findings Using Secondary Dataset Results from the Oh *et al* dataset, which was analyzed using the same workflow as the primary dataset. **A)** Scatter plot depicting the candidate contigs considered for analysis in this study. Each point is a contig that mapped to a reference virus genome. The x-axis shows the length (in nucleotides) of the contig subsection that mapped to the reference genome. The y-axis shows the overall coverage of the contig as a quantification of sequences aligning to the contig. The color highlights the reference virus genome that the contig was most similar to, and the size depicts the blast bit score associated with the contig-reference match. **B)** Box plots depicting the evolutionary pressure of *Staphylococcus* bacteriophages at the hypervariable loci (blue) and the regions immediately adjacent to the hypervariable loci (red). **C)** Heat map portraying the counts of every possible nucleotide substitution for each SNP found within *Staphylococcus* phage hypervariable loci. Tile color weight corresponds to the relative abundance of SNP substitution counts. The diagonal line highlights the panels associated with no substitution. The substitution patterns of amino acids at each **(D)** SNP, **(E)** amino acid charge, and **(F)** polarity with acidity are also shown. **G)** Notched boxplot illustrating the percent of primary dataset SNPs whose nucleotide positions were identical to those from the secondary validation sample set (left) compared to a simulated dataset of randomly assigned SNP locations (right). The inset shows an example contig identified in both datasets with 81% identical SNP positions. SNPs are represented as yellow lines, with the inner circle representing the validation dataset, and the middle circle representing the primary dataset. The outmost ring illustrates the contig, colored by nucleotides (A=Red, C=Blue, G=Yellow, T=Green). Boxplot parameters as described in figure 1.

Similar to our primary analysis, we identified 158 hypervariable loci within the *Staphylococcus* phage communities, and observed only 12 hypervariable loci associated with *Propionibacterium* phages, further highlighting the overall lack of genetic variability of the *Propionibacterium* phage communities. The *Staphylococcus* phage hypervariable loci were associated with purifying selection, yielding a ***pN**/**pS*** ratio slightly below 0.4 (**Figure 5B**), recapitulating our findings in the primary dataset (**Figure 1E**).

*Staphylococcus* phage nucleotide substitutions were associated with transitions between guanine and adenine residues, as we observed in our primary analysis (**Figure 5C**). The ti/tv ratio of these loci was 1.02. The most common amino acid substitution was proline to leucine (**Figure 5D**) and the substitution properties appeared loosely specific based on charge and polarity (**Figure 5E-F**), reproducing our findings (**Figure 2**).

We also evaluated the reproducibility of SNP position between identical but independently assembled genomic contigs of the two studies. We quantified the proportion of SNPs in our primary dataset that were also found at the same position in the secondary dataset. This revealed a median of approximately 50% overlap between datasets (**Figure 5G**). As a control, we generated a simulated dataset using randomly assigned SNP positions instead of those determined experimentally. This yielded a significantly lower median of approximately 15% shared nucleotide SNP calls (**Figure 5G**), suggesting that the observed SNP position is not random. These data indicate that our findings are consistent across different skin virome populations and techniques of collection and sequencing.

## DISCUSSION

Here we report localized targeted hypervariability in some of the most prevalent members of the skin virome. Hypervariable loci provide a substrate for complex virus evolution throughout the virome, which manifest as natural selection that differs by virus type and enforces purifying selection. Hypervariable loci, which were present in genes encoding factors for virus tropism and host immune evasion, were under purifying selection, whereas overall virus genomes were under near neutral selection. We characterized selected substitution of nucleotides within hypervariable loci, with different variant patterns between HPVs and *Staphylococcus* bacteriophage communities. These findings were validated in an independently collected cohort.

We showed *Propionibacterium* phages exhibited strikingly low nucleotide variation with nearly no identifiable hypervariable loci. While this starkly contrasts with HPVs and *Staphylococcus* bacteriophages, it agrees with our current understanding of *Propionibacterium* phage diversity. Genome comparisons of *Propionibacterium* phage isolates revealed minimal nucleotide diversity, although this has yet to be supported by targeted metagenomic evidence such as what we are presenting here (Marinelli et al., 2012; Liu et al., 2015). The lack of *Propionibacterium* phage hypervariability in our metagenomic dataset provides another level of evidence for minimal *Propionibacterium* phage diversity on the skin.

There are many potential factors that could contribute to the limited diversity of *Propionibacterium* phages, and a consensus has yet to be reached. The lack of hypervariable loci suggests minimal evolutionary pressure on the phages, which may likely be a reflection of their environment. As has been suggested previously, the phages and their hosts reside in a unique and relatively isolated environment deeper in the skin, which may contribute to low genomic diversity (Marinelli et al., 2012). Our data further support this hypothesis. More studies will be required to better understand this striking lack of genomic diversity observed in *Propionibacterium* bacteriophages.

We observed greater selective pressure on HPVs compared to *Staphylococcus* phages, which may reflect greater pressures from the human immune system, compared to phage bacterial hosts. This may also reflect the effects of different virus replication-cycles on evolutionary selection. HPVs do not usually exist in a latent, integrated state, while *Staphylococcus* phages do (Bae et al., 2006; Goerke et al., 2009; Edwards et al., 2013). As long as the *Staphylococcus* phages are integrated into the bacterial genome, we hypothesize that they are under less selective pressure by external factors.

The viral hypervariable loci are primarily associated with purifying selective pressure, a finding in agreement with previous non-metagenomic virus reports (Chen et al., 2005; Wolf et al., 2006; Li et al., 2011). The observed prominent purifying selective pressure supports an evolutionary model of long static periods punctuated by brief positive selection, as is observed in influenza virus (Wolf et al., 2006). Here nucleotide diversity acts as a primer for rapid virus adaptation through brief positive selection, while maintaining periods of consistency through purifying selection during static environmental conditions. Longitudinal studies will be required to further support this model.

The amino acid substitutions associated with hypervariable loci were non-random, and followed virus-specific substitution patterns (**Figure 2**). HPV hypervariable loci were most associated with substitutions from glycine to valine. This substitution has recently been associated with infectious functionality, whereby the introduction of this mutation resulted in impaired infective ability of the virus (Bronnimann et al., 2013). This impaired infectious activity was attributed to a reduced efficacy of genomic DNA endosomal translocation within the host, which may have been the result of impaired trans-membrane alpha-helical self-association of the L2 minor capsid protein. Given these findings, our results suggest hypervariable loci are involved in promoting diversity in endosomal translocation motifs to some degree. Hypervariable loci may certainly have other diverse, functional roles, as evidenced by the wide range of hypervariable loci-containing genes.

The dominant amino acid substitution observed in *Staphylococcus* bacteriophages was from proline to leucine, a different substitution than that observed in HPV. This substitution could affect protein structure, particularly a loss of rigidity due to the loss of the proline ring structure. This observation may reflect a biologically important adaptation of the bacteriophage to its *Staphylococcus* host, which have been shown to be auxotrophic for proline and leucine and may switch between auxotroph and prototroph depending on nutrient availability (Emmett & Kloos, 1975; Nuxoll et al., 2012). Because the amino acids may be in variable supply depending on the host, phages may alter their amino acid usage to exploit what is most readily available.

The overall selective nucleotide substitutions associated with HPV amino acid charge highlights a potential maintenance of HPV tropism. The lack of HPV substitutions between charges may suggest a selection against strong alterations in protein isoelectric points, which have been implicated in affecting HPV tropism (Mistry, Wibom & Evander, 2008). Furthermore, because acidic residues almost never mutated to non-polar residues, these acidic amino acids are potentially important external amino acids that may participate in tropic protein-protein interactions.

The described patterns in our findings suggest a role for targeted and/or localized genomic variation. One mechanism of such active targeted variation in *Staphylococcus* bacteriophages is diversity generating retroelements. In this system, a phage-encoded reverse transcriptase copies a template region to create a variable region in a gene in an error-prone fashion. We identified such an element that is likely active and promotes diversity in a gene of unknown function. While informative, this discovery only explains the diversity-generating mechanism of a small proportion of hypervariable loci. Significant further investigation will be needed to characterize other potential mechanisms.

This study illustrates the diversity of evolutionary pressures on skin virus communities. It begins to provide further community-wide context to the molecular understanding of skin viruses, and highlights important aspects of their infectious cycles. These insights also contribute to understanding virus ecology of the human skin, and will inform future translational research into HPV vaccination, vaccination against other skin-associated viruses, effects of phages on bacterial pathogenesis, and phage therapy. Understanding how viruses evolve in their natural communities is crucial for improving these translational applications, and our findings here, which focus on HPV and *Staphylococcus* phages, will benefit cutaneous clinical virology and provide a foundation for future studies.

## CONCLUSIONS

We report that the skin virus communities contain hypervariable loci that are associated with strong purifying selection and targeted nucleotide substitution. The degree of selective pressure and impact of amino acid substitutions on protein chemistry (structure, isoelectric point, polarity) is virus specific, despite being members of the same community. These hypervariable loci are found within diverse viral strains, with varying degrees of phylogenetic divergence over their evolutionary history. We further reproduce these findings in independently collected skin virus communities.

## ACKNOWLEDGEMENTS

We thank the members of the Grice and Bushman laboratories for their underlying contributions.

## FORMULAS

Formula 1

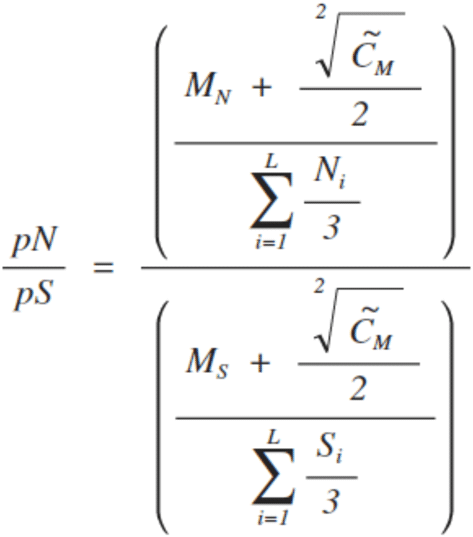

The formula used to calculate the ***pN/pS*** ratio for a gene. M_N_ is the number of non-synonymous SNPs within the gene and M_S_ is the number of synonymous SNPs found within the gene. Each mutation value is normalized for the likelihood that the result would have happened by chance, calculated as the sum of the proportions of nucleotides that would have resulted in either a non-synonymous or synonymous mutation. To calculate this proportion, the possible non-synonymous mutations (N) and synonymous mutations (S) at position i within the gene are expressed as a fraction of the three possible alternate nucleotides. SNP counts were smoothed as pseudo-counts using the median SNP sequence coverage (C_M_).

## SUPPLEMENTAL FIGURES

**Figure S1.**
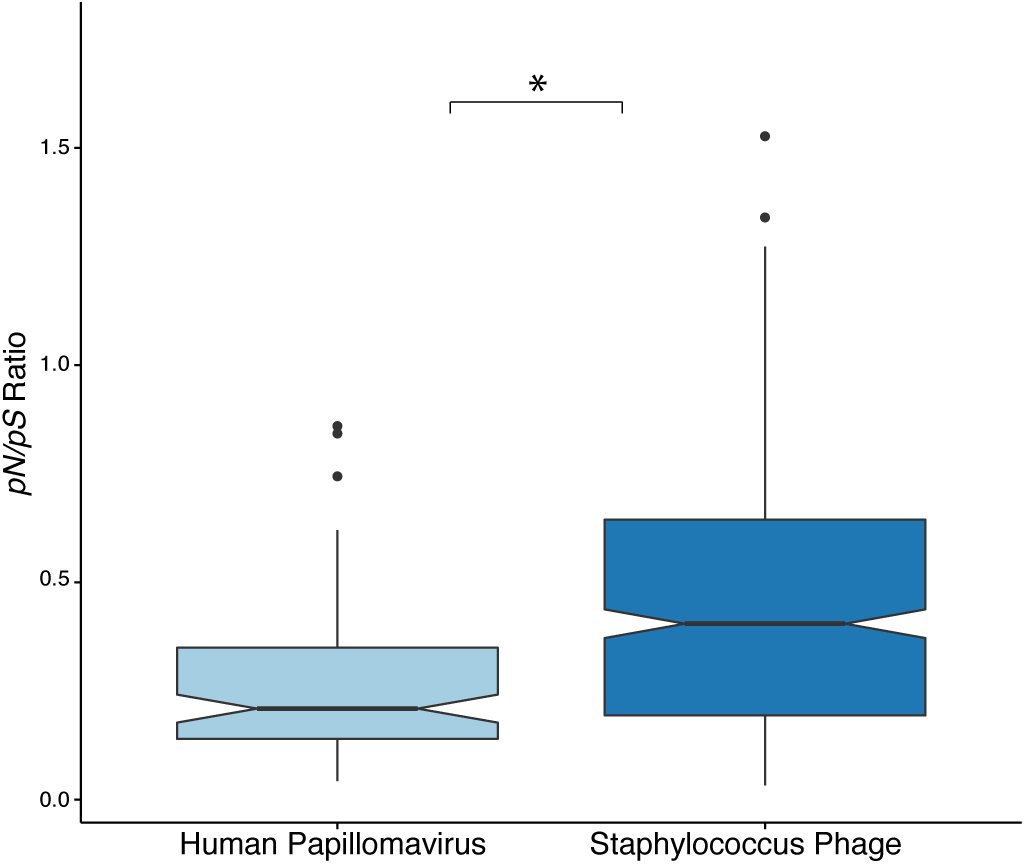
Selective Pressure of Hypervariable Loci Between Viruses Notched box plot of the hypervariable loci selective pressures between HPV and *Staphylococcus* phages. The difference between these populations is significant by Wilcoxon test (p < 0.005; marked by asterisk).

**Figure S2.**
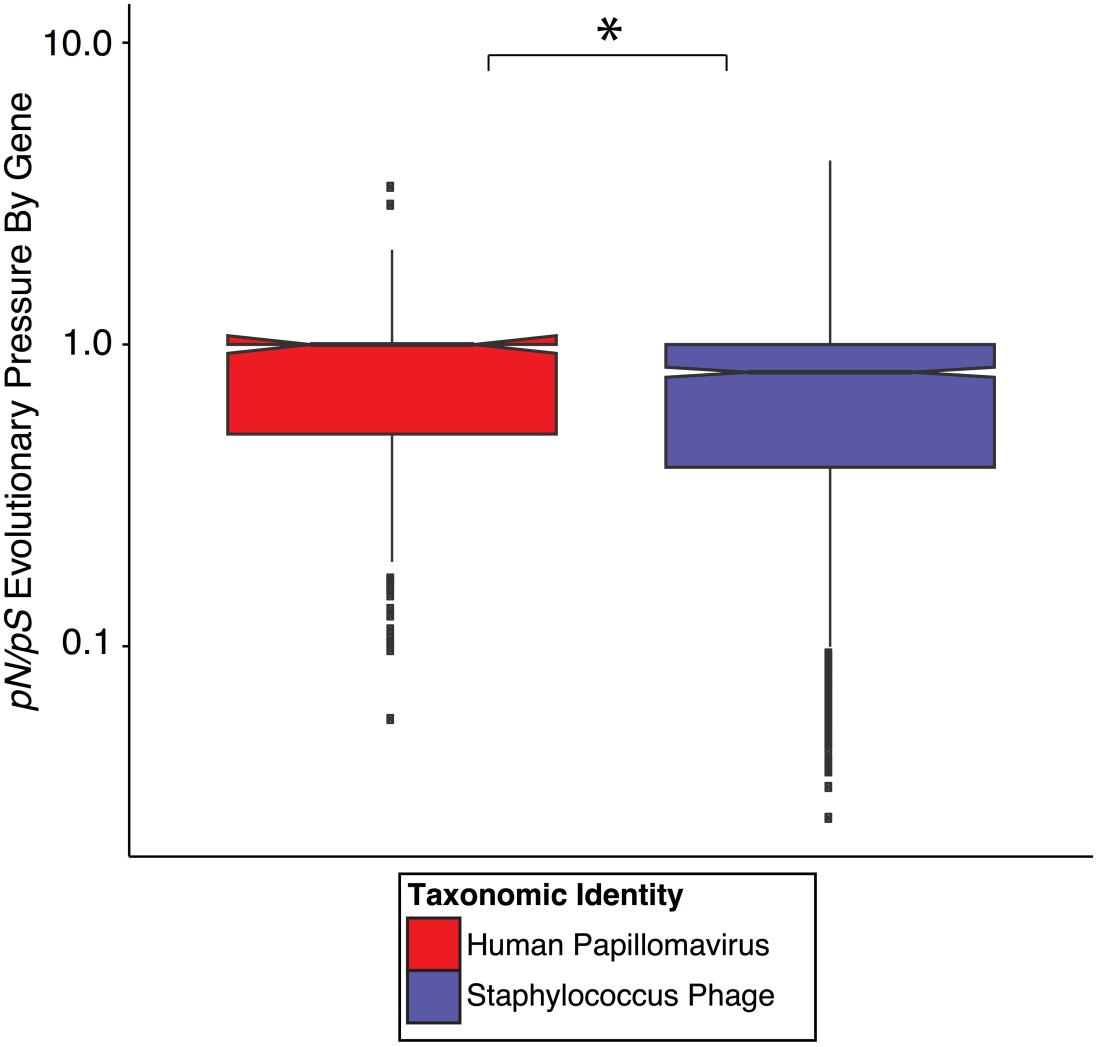
Overall Selective Pressure of Virus Genomes Box plot of the comprehensive evolutionary pressure on each virus of interest. The y-axis depicts the ***pN/pS*** ratio of genes from each virus denoted on the x-axis. Viruses are highlighted in red (HPV) and blue (*Staphylococcus* phage). The difference between these populations is significant by Wilcoxon test (p < 0.01; marked by asterisk).

